# Impact of Chrysin on Vitamin D and Bone Health - Preclinical Studies

**DOI:** 10.1101/2020.11.19.390757

**Authors:** Siva Swapna Kasarla, Sujatha Dodoala, Sunitha Sampathi, Narendra Kumar Talluri

**Affiliations:** Institute of Pharmaceutical Technology, Sri Padmavati Mahila Viswavidyalayam, Tirupati, India 517502; GITAM school of Pharmacy, GITAM (Deemed to be University), Sangareddy, Hyderabad, India 502329; Department of Pharmaceutical Analysis, National Institute of Pharmaceutical sciences (NIPER), Balanagar, Hyderabad, India 500037

**Keywords:** Bone, Chrysin, CYP3A4, Vitamin D deficiency, LC-MS/MS, Osteoporosis, 25-OH-D_3_

## Abstract

Vitamin D deficiency is an endemic problem existing worldwide. Although several strategies were established to enhance vitamin D_3_ levels, studies specifically focussing inhibition of vitamin D metabolism which may prolong the availability of active vitamin D in pathological conditions are less explored. Studies also suggest that higher doses of vitamin D_3_ fail to achieve optimum vitamin D levels. In this context, we focussed on the enzyme CYP3A4 which promotes inactivation of active vitamin D. The current study was aimed to decipher the impact of chrysin, a proven CYP3A4 inhibitor as an intervention and its effects in combination with low dose vitamin D_3_ (40 IU) and bone health in vitamin D deficiency condition. The *in-vivo* activity of chrysin was evaluated on female Wistar albino rats fed with a vitamin D deficient diet to attain vitamin D deficiency for 28 days. Chrysin was given alone and in combination with calcium carbonate (CaCO_3_) and/or vitamin D_3_. All the therapeutic interventions were assessed for serum 25-OH-D3 by LC-MS, biochemical, urinary, and bone parameters. Animals treated with chrysin alone and in combination with low dose vitamin D_3_ and/or CaCO_3_ showed an eminent rise in serum 25-OH-D3 levels along with increased serum biochemical parameters. On contrary, a significant decrease in the urinary parameters followed by beneficial effects on bone parameters was noticed in contrast with the vitamin D deficient diet group. Our findings revealed that although chrysin alone showed a notable effect on 25-OH-D3 and osseous tissue, comparatively it showed intensified therapeutic effect in combination with vitamin D_3_ and CaCO_3_ which can be employed as a cost-effective option to improve bone health.

**Graphical Abstract:** 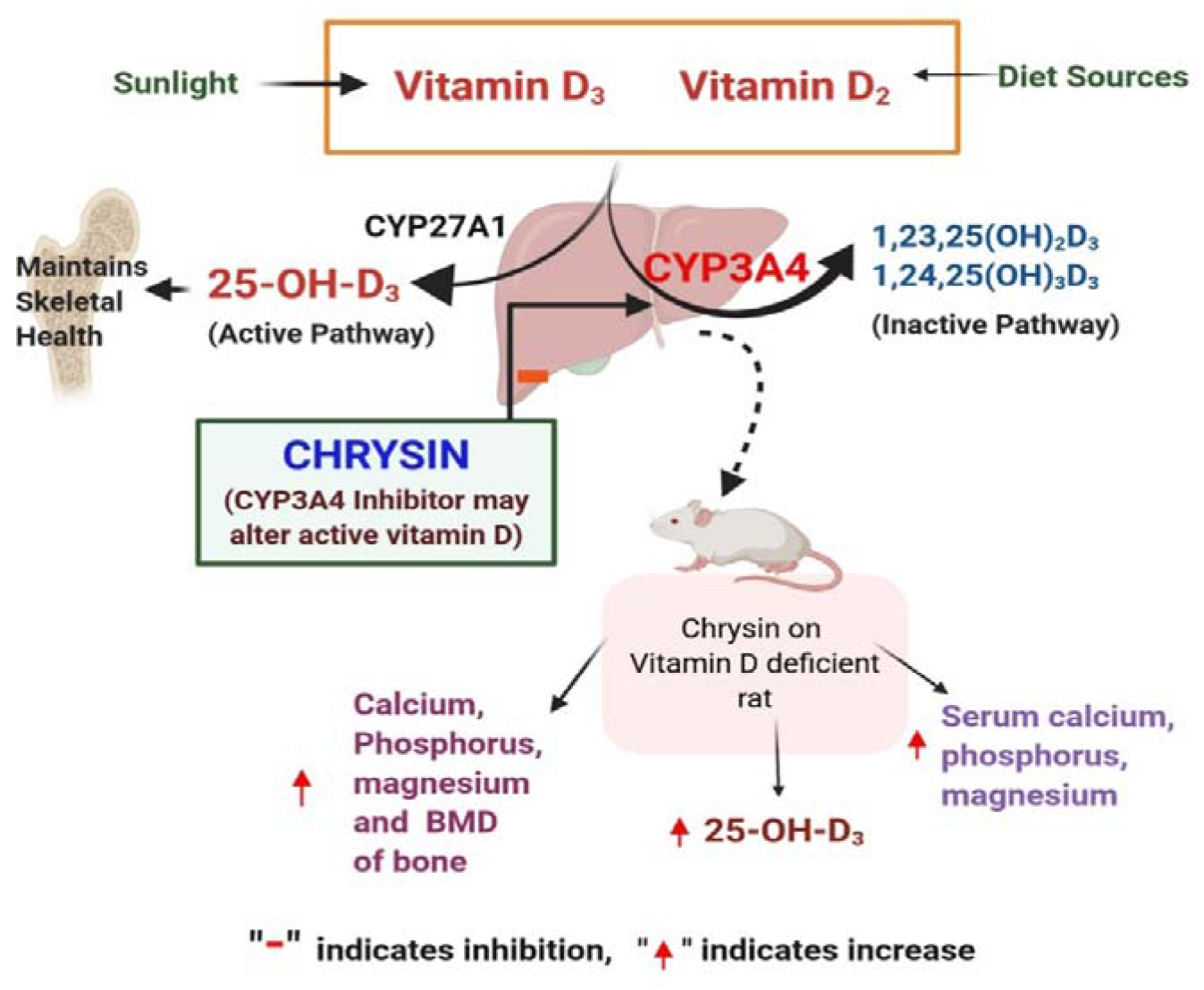

## Introduction

Currently, the magnitude of musculoskeletal diseases is emerging systematically among all the age groups affecting the social life of the people leading to pain and disability [1]. In particular, osteoporosis is the most discussed burden of about 1 in 3 women and 1 in 5 men [2]. The predominant risk factors projecting bone diseases like osteoporosis are age, sex, genetic defects, sedentary lifestyle, vitamin D deficiency/insufficiency, minimal intake of calcium and early menopause [3]. However, the possible aspects that can reduce the intensity of skeletal disorders can be recognition of the disease in its early stage followed by effective treatment which includes ingestion of appropriate vitamin D and calcium supplements to combat bone loss [4].

Vitamin D is broadly recognized as a “bone vitamin” for its magnificent impact on osseous tissue. Conversely, its insufficient distribution leads to vitamin D deficiency condition which has a tremendous effect on skeletal health and in worsening the pathological conditions like osteoporosis [5–6]. It was formerly known that the imperative relationship between active vitamin D and transcellular calcium uptake is necessary for maintaining bone architecture [7]. The observational findings on calcium kinetics specified that systematic absorption of calcium is about 30-40% in case of adequate vitamin D (>32ng/ml) and elicits only 10-15% in case of low vitamin D (<15ng/ml) [8].

On the other hand, though the hypovitaminosis D is a modifiable risk factor, the medical community is facing several challenges in terms of maintaining or achieving optimum serum vitamin D_3_ levels and retaining bone mass [9]. Respective research strategies are implementing to conquer vitamin D insufficiency targeting sunlight exposure and climatic conditions [10], gut absorption of vitamin D supplements [11], vitamin D receptor (VDR) genotyping [12], metabolic expression of cytochrome P450 (CYP450) enzymes and their interplay with vitamin D [13,14]. CYP450 enzymes are the major class of catabolic enzymes which play a decisive role in pharmacokinetics of drugs as well as endogenous substances. As 90% of therapeutic agents are metabolized of CYP enzymes, the multitude of studies implying CYP450 gained outmost importance to unveil various unpredicted drug-drug interactions [15]. Possibly the interactions caused by enzymatic inhibition/induction results in diminished/enhanced metabolic compounds which may have greater emphasis on kinetic and dynamic profiles of therapeutic targets [16].

In this regard, we focussed on the multifunctional class of enzyme CYP3A4 which catalyses 25-hydroxylated mediated conversion of the active form of vitamin D (1,25(OH)_2_D_3_) into its inactive form [14]. Interestingly, research studies revealed that therapeutic substrates (Inducers/Inhibitors) of CYP3A4 enzymes may contribute to alter the 1,25(OH)_2_D_3_ levels in the serum [15]. In this perspective, subsequent inhibition of vitamin D metabolism by CYP3A4 substrates may show ameliorative effect on hypovitaminosis D. A systematic study stated that higher doses of vitamin D_3_ fail to achieve optimum serum 25-OH-D_3_ levels in humans [17]. So, we gave a thought to enhance vitamin D levels by incorporating a well-established CYP3A4 inhibitor chrysin, a natural flavonoid having beneficial pharmacological effects such as anti-cancer, neuroprotective, hepatoprotective, cardioprotective, anti-arthritic effect [18, 19]. It has also shown a promising outcome in osteoporotic rats [20]. Our research aimed to study the impact of chrysin on low dose vitamin D_3_ supplementation and its further implications on vitamin D deficiency condition and bone health.

## Materials and Methods

### Materials

Chrysin, also known as 5,7-dihydroxyflavone acquired from Sigma Aldrich, India. Vitamin D_3_ capsules of 2000 IU were used and obtained from UAV Private Limited, Mumbai marketed as D-RISE 2000 soft gelatin capsules. Calcium carbonate and all other chemicals used for the preparation of vitamin D deficient diet were of analytical grade and purchased from SD-fine chemicals and Merk Pvt Ltd, India. The kits used for estimation of various biochemical parameters were from Erba Manheim, Pvt Ltd.

### Preparation of vitamin - D deficient diet

The required ingredients for the preparation of the diet were weighed accordingly and blended for about 45 minutes followed by addition of 10% water in order make a dough. Then it is granulated by using the sieve 8mm followed by making round shaped balls, dehydrated and dried for about 3 days at room temperature of 37ΰ C. The diet prepared was standardized to ensure the uniform nutrient values as that of normal diet. The prepared diets were stored at 2-8ΰC to minimize oxidation [21]. The composition of the normocalcemic vitamin D deficient was followed as per the earlier reference [22] [Suppl. Table 1]

### Selection of doses

Based on the earlier research studies the doses of chrysin, vitamin D_3_, CaCO_3_ were fixed at 100 mg/kg, 40 IU, 50 mg/kg respectively [23,24].

### Experimental animals

Adult healthy female Wistar albino rats weighing 180-200g were obtained from a certified breeder Sri Venkateshwara Enterprises, Bangalore, India and housed under standard environmental conditions like ambient temperature (22 ± 2ΰC), relative humidity (30-70%) and a 12/12hr light/dark cycle in individual polypropylene cages and maintained hygienic conditions throughout the study according to the committee for the purpose of control and supervision of experiments on animals (CPCSEA) guidelines. All experimental protocols were approved by the Institutional Animal Ethics Committee [Approval. No. CPCSEA/1677/PO/Re/S/2012/IAEC/34].

### *In vivo* experimental design

Vitamin D deficiency was induced by the diet prepared. According to the research of interest the animals were divided into six groups (n=6) and all the treatment groups (group II, III, IV, V, VI) were fed with vitamin D deficient diet except normal (group I) followed by treatment for 28 days and the schedule was as follows.

Group I: Normal (standard laboratory diet and drinking water *ad libitium*)

Group II: Disease control (fed with vitamin D deficient diet)

Group III: Therapeutic intervention 1 (chrysin (100 mg/kg, *po*))

Group IV: Therapeutic intervention 2 (chrysin (100 mg/kg, *po*) + CaCO_3_ (50 mg/kg, *po*))

Group V: Therapeutic intervention 3 (chrysin (100 mg/kg, *po*) + vitamin D_3_ (40 IU/kg, *po*))

Group VI: Therapeutic intervention 4 (chrysin (100 mg/kg, *po*) + CaCO_3_ (50 mg/kg, *po*) + vitamin D_3_ (40 IU/kg, *po*))

The required quantities of test drug solution (chrysin), calcium carbonate (CaCO_3_) was prepared by suspending in 1% w/v carboxy methyl cellulose (CMC) and vitamin D_3_ in distilled water on every day before administering to the animals. The prepared solution was administered with the help of an oral feeding needle (18G) accordingly.

### Body weights

The body weights of all the animals were taken at weekly intervals i.e., on 7^th^ and 28^th^ day of the study and compared the difference in the weights of animals under different groups.

### Biochemical evaluation

The blood samples were collected from the retro-orbital plexuses at regular intervals i.e., on 7^th^ and 28^th^ day of the study. The serum was separated and analysed for calcium, phosphorus, magnesium, and alkaline phosphatases with the help of kits by using autoanalyzer (Erba Mannehim) respectively. The collected serum samples were also used for the estimation of vitamin D metabolites for which the samples were processed and stored in amber coloured eppendorf tubes at 4°C, to avoid degradation of vitamin D metabolites [25].

### Estimation of urinary parameters

The animals were hydrated with 5ml of water and were individually housed in the metabolic cages for 6 hrs and the collected urine was then centrifuged at 2000 rpm for 5 minutes. The supernatant collected and stored in air-tight containers at 4-8ΰC for further evaluation of urinary excretion of calcium, phosphorus and magnesium.

### Estimation of 25-OH-D_3_ by LC-MS/MS

The concentration of 25-OH-D_3_ in the serum was quantified by using most reliable LC-MS/MS method. The extraction of 25-OH-D_3_ from the serum samples was obtained by simple liquid-liquid extraction. Initially, 200 μl of serum sample was mixed with 25 μl of internal standard in 200 μl of methanol and equilibrate at 5 minutes. Then 200 μl of methanol was added and vortex for 10 seconds and let stand for precipitation. Then 2 ml of n-heptane was added, and vortex for 10 minutes. The samples were centrifuged at 10,000 rpm for 10 minutes to separate organic layer. The organic layer was transferred into clean tubes and evaporated to dryness under nitrogen at room temperature. The prepared samples were reconstituted in 100 μl of 2 mM ammonium acetate, 0.1% formic acid in 80:20 methanol/water [26, 27].

The 20 μl of the sample is initially analysed using an Agilent 1200 series HPLC instrument (Agilent Technologies USA) with C18 (250 mm x 4.6 mm i.d.) column maintained at ambient temperature (30ΰC) coupled to a quadrupole time-of-flight (Q-TOF) mass spectrometer (Q-TOF-LC/MS 6510 classic G6150A, Agilent technologies, USA) with a dual ESI as ion source. The data acquisition was under the control of Mass Hunter workstation software. The mobile phase A: 2 mM ammonium acetate, 0.1% formic acid in water, B: 2 mM ammonium acetate, 0.1% formic acid in methanol was filtered with 0.45μm nylon membrane and pumped with a flow rate of about 10 mL/min. The standard calibration curve of 25-OH-D_3_ was plotted by selected serial dilutions of 5,10,15,20,25 ng/ml. The standard 25-OH-D_3_ peak was identified and quantified by comparing the selected fragment ion (m/z 383.5→401.3).

### Bone parameters

On the day 28^th^ day of the study the animals were sacrificed by cervical dislocation. The bones such as femur, tibia – fibula, 4^th^ lumbar and 8^th^ thoracic vertebrae were collected, weighed and stored. The stored bone samples were further used to carry out various bone parameters as follows

#### a. Bone weight

Freshly isolated femurs and tibia fibula were taken, and adhering tissues were removed and weighed by using electronic digital balance [28]

#### b. Bone length

The isolated femurs and tibia fibula were cleaned by removing adhering tissues and length is measured from proximal tip of the head to the distal tip of medial condyle by using vernier callipers scale [28].

#### c. Bone hardness

The isolated femurs and tibia fibula were cleaned and was placed in Pfizer hardness compression tester and the pressure was applied till the bone breaks. The breaking strength is considered as bone hardness and the readings were obtained in Newton’s [29].

#### d. Vertebral hardness

The fourth lumbar vertebra and eighth lumbar vertebra was spotted, isolated and were placed in Pfizer hardness compression tester and the readings were obtained in Newton’s [29].

#### e. Bone mineral density

The bones were cleaned and freed of soft tissue and then frozen for further analysis of bone mineral density. Before examination bones were defrosted for about 30 min followed by analysis using dual energy X – ray absorptiometry (Elite ACCLAM series) for a scan time of about 3 min and results obtained were evaluated accordingly [30,31].

#### f. Bone ash content

The bones were cleaned and adhering soft tissue is removed for the estimation of bone mineral content. They were kept in muffle furnace at 600ΰC for 24hrs. The collected bone ash was weighed with the help of digital balance and further diluted with 6 N HCl and assayed for calcium, magnesium, and phosphorus content by calorimetric estimation using respective diagnostic kits [31].

### Histopathology

Bone specimens collected from various treatment groups were fixed in 10% formalin followed by further decalcified and dehydrated with ascending grade of alcohol and washed with distilled water and were processed for paraffin sectioning. Finally, the sections were stained with haematoxylin and eosin for evaluation. The prepared sections were focused under fluorescence Microscope (Olympus Private Limited) for histopathological changes [32].

### Statistical Analysis

Data was expressed as mean ± standard deviation (SD) of 6 observations, and evaluated by using one – way and/or two – way analysis of variance (ANOVA) followed by Tukey’s multiple comparison test by using graph pad prism software version 8 (Graph pad software, Inc. La Jola, CA, USA). The confidence level of p ◻0.05 was statistically significant.

## Results

### Effect of therapeutic interventions on body weight (gm)

A significant (p <0.05) decrease in the body weight of animals treated with vitamin D deficient diet (group II) was noticed at regular intervals, compared with normal animals (group I). In case of animals treated with chrysin and in combination with CaCO_3_ and/or Vitamin D_3_ (group III, IV, V, VII) have shown a significant (p ◻0.05) increase in the body weight when compared to diet control group indicated the positive impact to improve vitamin D deficiency. The overall effect of drug treatments on body weight were summarized in Table 1.

**Table 1:**
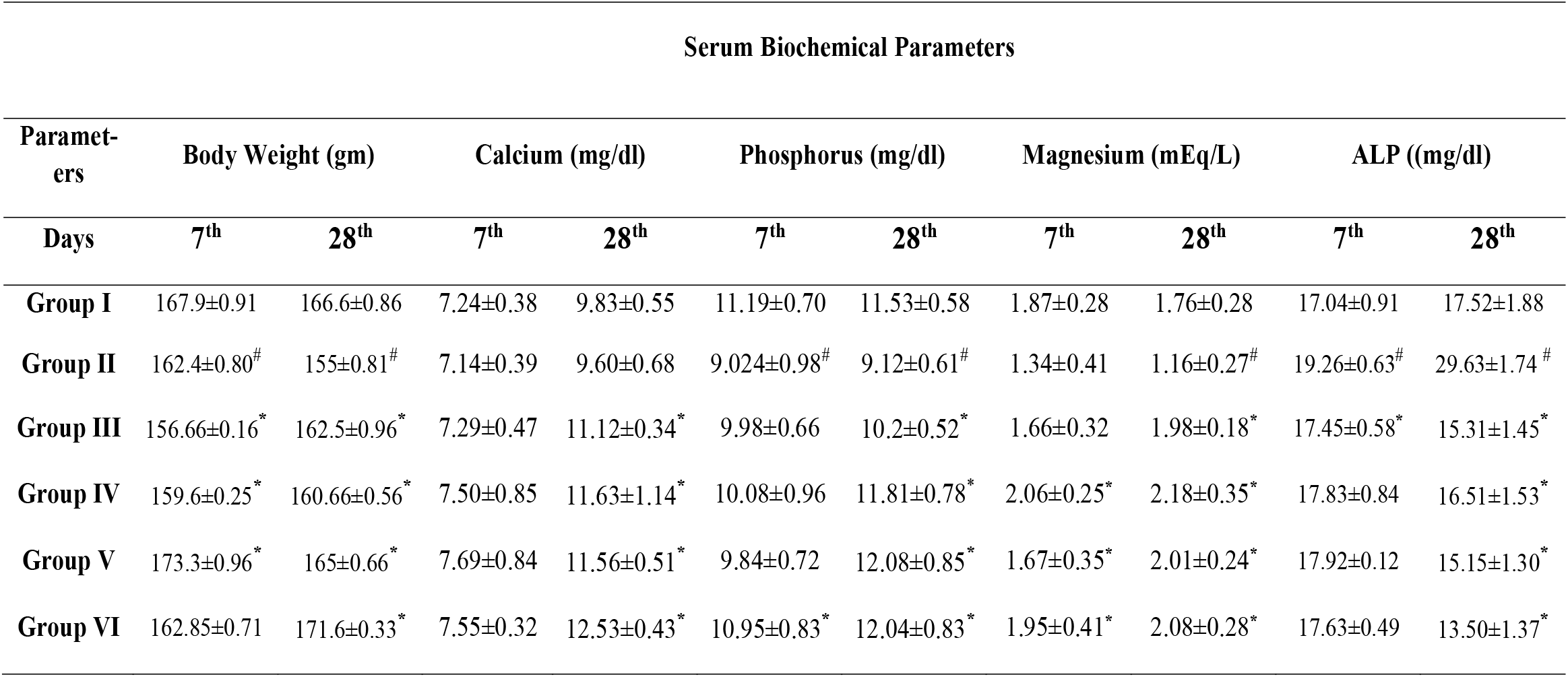
Effect of therapeutic interventions on serum calcium (mg/dl), phosphorus (mg/dl), magnesium (mEq/L), and ALP ((mg/dl) levels. [data were expressed as mean ± SD (n=6); analysed by two-way ANOVA followed by Tukey’s multiple comparison test. # = p<0.05, considered statistically significant when compared group I (normal) vs group II (diet control group). * = p<0.05, considered statistically significant when compared to the other treatment groups (group III, IV, V, VI) vs. diet control group]

### Effect of therapeutic interventions on serum calcium (mg/dl), phosphorus (mg/dl) levels, magnesium (mEq/L) levels

Research study revealed that there is no significant effect on serum calcium levels of disease control animals (group II) with respect to normal animals which resembles the diet supplied doesn’t modulate calcium levels. Whilst there is a significant decrease in the serum phosphorus, magnesium levels effect among DC animals with respect to normal animals which particularly presents the significance of vitamin D_3_ in mineral homeostasis. Whereas other treatment groups such as III, IV, V, VI showed a significant (p ◻0.05) increase in serum calcium, phosphorus and magnesium at regular intervals, compared to the vitamin D deficient diet control group. This suggests the definite effect of chrysin in vitamin D deficient condition and were illustrated in table 1.

### Effect of therapeutic interventions on serum ALP (mg/dl) levels

A significant (p ◻0.05) increase in the serum ALP levels was noticed among animals in group received vitamin D deficient diet, compared to the group I which indicates bone deterioration. But, a significant (p ◻0.05) decrease in the serum ALP levels was noticed in the groups III, IV, V, VI, when compared to group II animals fed with vitamin D deficient diet, which revealed the decreased bone damage in contrast with disease control (DC) animals (Table 1).

### Effect of therapeutic interventions on urinary calcium (mg/dl) levels, magnesium (mEq/L) levels, phosphorus (mg/dl) levels

Study showed a significant (p ◻0.05) surge in excretion of urinary calcium, magnesium, phosphorus levels in vitamin D deficient diet treated animals, when compared to normal animals. This hinted towards the increased excretion of calcium, magnesium, phosphorus which coincides the decreased availability in the serum. A significant (p ◻0.05) decline in the excretion of urinary parameters were found in the groups treated with chrysin along with other treatment combinations i.e., groups III, IV, V, VI when compared to vitamin D deficient diet groups (Table 2).

**Table 2:** Effect of therapeutic interventions on urinary calcium (mg/dl) levels, magnesium (mEq/L) levels, phosphorus (mg/dl) levels. [data were expressed as mean ± SD (n=6); analysed by two-way ANOVA followed by Tukey’s multiple comparison test. # = p<0.05, considered statistically significant when compared group I vs group II. * = p<0.05, considered statistically significant when compared to the other treatment groups (group III, IV, V, VI) vs. diet control group]

| Urinary Parameters |  |  |  |  |  |  |
| --- | --- | --- | --- | --- | --- | --- |
| Parameters | Calcium (mg/dl) |  | Phosphorus (mg/dl) |  | Magnesium (mEq/L) |  |
| Days | 7<sup>th</sup> | 28<sup>th</sup> | 7<sup>th</sup> | 28<sup>th</sup> | 7<sup>th</sup> | 28<sup>th</sup> |
| Group I | 6.68±0.49 | 6.98±0.40 | 1.21±0.02 | 2.91±0.22 | 1.4±0.06 | 2.45±0.022 |
| Group II | 7.73±0.59<sup>#</sup> | 10.4±0.38<sup>#</sup> | 1.03±0.021 | 4.61±0.19<sup>#</sup> | 1.98±0.098<sup>#</sup> | 4.88±0.98<sup>#</sup> |
| Group III | 6.88±0.30<sup>*</sup> | 6.05±0.22<sup>*</sup> | 0.88±0.01 | 2.31±0.12<sup>*</sup> | 0.92±0.021<sup>*</sup> | 2.94±1.20<sup>*</sup> |
| Group IV | 6.76±0.73<sup>*</sup> | 5.11±0.42<sup>*</sup> | 1.63±0.09 | 1.97±0.12<sup>*</sup> | 1.02±0.065<sup>*</sup> | 1.99±0.06<sup>*</sup> |
| Group V | 6.165±0.67<sup>*</sup> | 5±0.41<sup>*</sup> | 1.92±0.06 | 2.01±0.11<sup>*</sup> | 1.24±0.08 | 2.42±0.071<sup>*</sup> |
| Group VI | 6.66±0.39<sup>*</sup> | 4.6±0.23<sup>*</sup> | 1.52±0.087 | 2.55±0.24<sup>*</sup> | 1.11±0.056<sup>*</sup> | 3.01±0.06<sup>*</sup> |

### Effect of therapeutic interventions on bone ash parameters such as ash weight (mg), calcium (mg/dl), phosphorus (mg/dl) and magnesium (mEq/L)

Among all the bone ash parameters, the disease control (group II) showed a significant (p ◻0.05) decrease in ash weight and calcium levels, which indicates the decrease in bone calcification and bone formation. Anyhow, no significant (p ◻0.05) difference in phosphorus, magnesium levels in bone ash samples of vitamin D deficient group was noticed, when compared to normal groups.

But, in case of animals treated with different therapeutic interventions such as groups III, IV, V, VI showed significant (p ◻0.05) increase in bone ash weight and calcium. This indicates, increased bone calcification and strong bone formation. A significant increase in the phosphorus levels of bone ash was noticed only in group 4 (Chrysin + CaCO_3_) and group 6 (Chrysin + CaCO_3_ + Vitamin D_3_) when compared to vitamin D deficient group. The levels of magnesium were not significantly altered between any of the groups (Table 3).

**Table 3:**
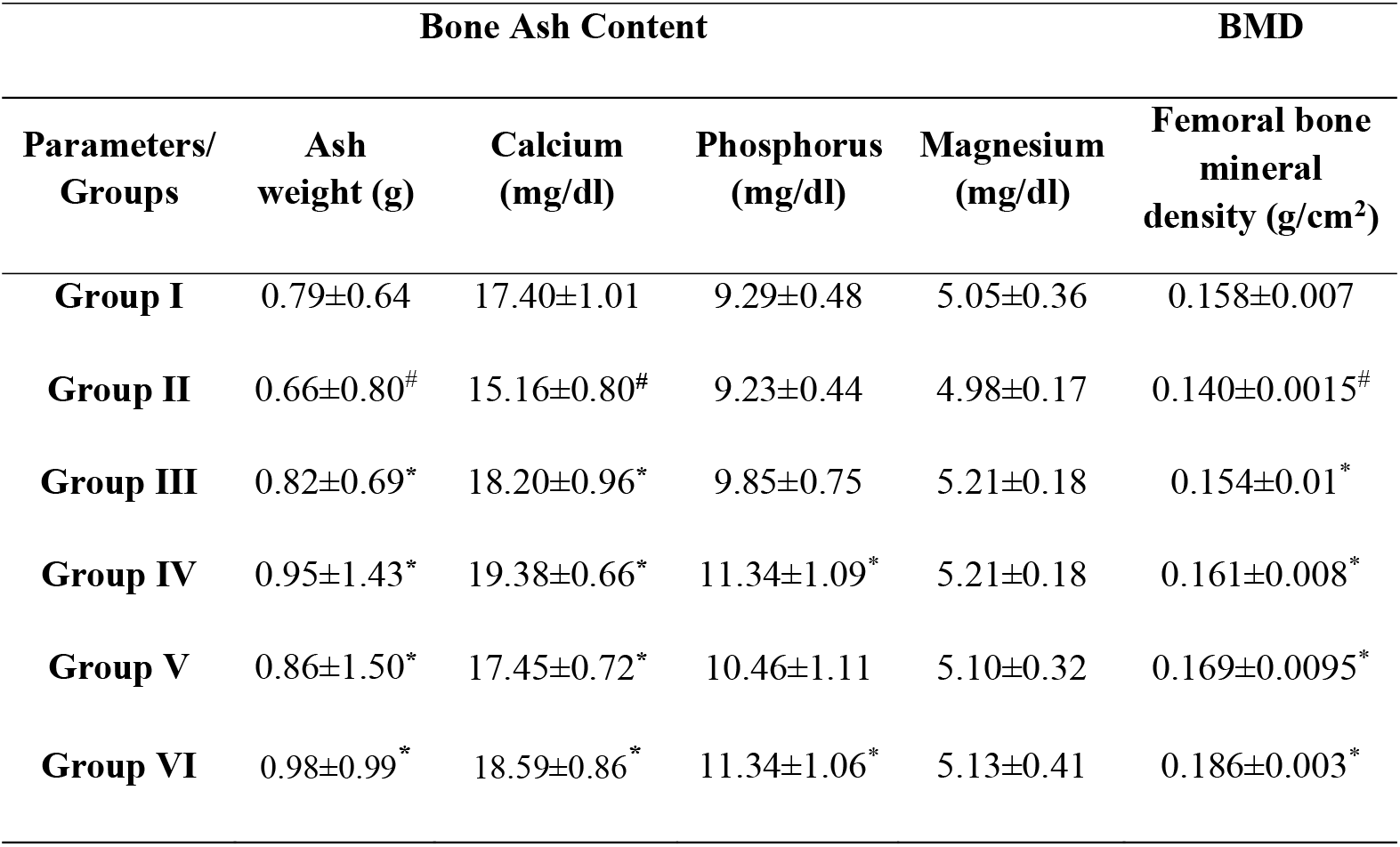
Effect of drug treatment groups on bone ash content (ash weight (mg), calcium (mg/dl), phosphorus (mg/dl) and magnesium (mEq/L) levels) and femur bone mineral density (g/cm^2^) [data were expressed as mean ± SD (n=6); analysed by one-way ANOVA followed by Tukey’s multiple comparison test. # = p<0.05, considered statistically significant when compared group I vs group II. * = p<0.05, considered statistically significant when compared to the other treatment groups (group III, IV, V, VI) vs. diet control group]

### Effect of therapeutic interventions on femur and tibia-fibula weights (gm), lengths (cm), and hardness’s (N)

Study showed that there is a significant (p ◻0.05) decrease in the femur and tibia-fibula weights and hardness’s in DC animals when compared to normal animals represents the decreased bone strength which may increase the risk of osteoporotic related fractures. In contrast, animals treated with chrysin and in combination with CaCO_3_ and/or Vitamin D_3_ showed significant (p ◻0.05) increase in the femur and tibia-fibula weights and hardness’s demonstrates the increased bone strength. However, no significant (p ◻0.05) difference was found in lengths of femur and tibia-fibula in any of the groups studied (Table 4).

**Table 4:** Effect of therapeutic interventions on femur and tibia-fibula weights (gm), lengths (cm), and hardness’s (N) [data were expressed as mean ± SD (n=6); analysed by one-way ANOVA followed by Tukey’s multiple comparison test. # = p<0.05, considered statistically significant when compared group I vs group II. * = p<0.05, considered statistically significant when compared to the other treatment groups (group III, IV, V, VI) vs. diet control group]

| Parameters/<br>Groups | FEMUR |  |  | TIBIA-FIBULA |  |  |
| --- | --- | --- | --- | --- | --- | --- |
|  | Length<br>(cm) | Weight<br>(gm) | Hardness<br>(N) | Length<br>(cm) | Weight<br>(gm) | Hardness<br>(N) |
| Group I | 2.93±0.10 | 0.74±0.06 | 9.8±0.21 | 3.27±0.23 | 0.72±0.036 | 14.17±2.44 |
| Group II | 2.93±0.19 | 0.61±0.08<sup>#</sup> | 8.94±0.12<sup>#</sup> | 3.23±0.27 | 0.55±0.07<sup>#</sup> | 6.98±1.91<sup>#</sup> |
| Group III | 3.02±0.10 | 0.74±0.03<sup>*</sup> | 9.93±0.24<sup>*</sup> | 3.55±0.27 | 0.66±0.035<sup>*</sup> | 11.91±2.10<sup>*</sup> |
| Group IV | 3.08±0.12 | 0.74±0.06<sup>*</sup> | 10.43±0.83<sup>*</sup> | 3.60±0.21 | 0.74±0.066<sup>*</sup> | 13.86±1.52<sup>*</sup> |
| Group V | 3.03±0.15 | 0.79±0.05<sup>*</sup> | 10.02±0.41<sup>*</sup> | 3.57±0.38 | 0.75±0.096<sup>*</sup> | 14.39±2.213<sup>*</sup> |
| Group VI | 3.13±0.12 | 0.88±0.06<sup>*</sup> | 11.76±0.67<sup>*</sup> | 3.65±0.33 | 0.84±0.016<sup>*</sup> | 15.60±2.70<sup>*</sup> |

### Effect of therapeutic interventions on 4th lumbar hardness (N) and 8th thoracic hardness (N)

Study showed that there was a significant decrease (p ◻0.05) in the 4^th^ lumbar hardness and 8^th^ thoracic hardness among group II animals compared to group I specifies the decreased bone mass or strength due to vitamin D insufficiency. But the animals treated with other therapeutic interventions showed a significant (p ◻0.05) increase in the 4^th^ lumbar hardness and 8^th^ thoracic hardness compared to animals fed with deficient diet (Table 5).

**Table 5:**
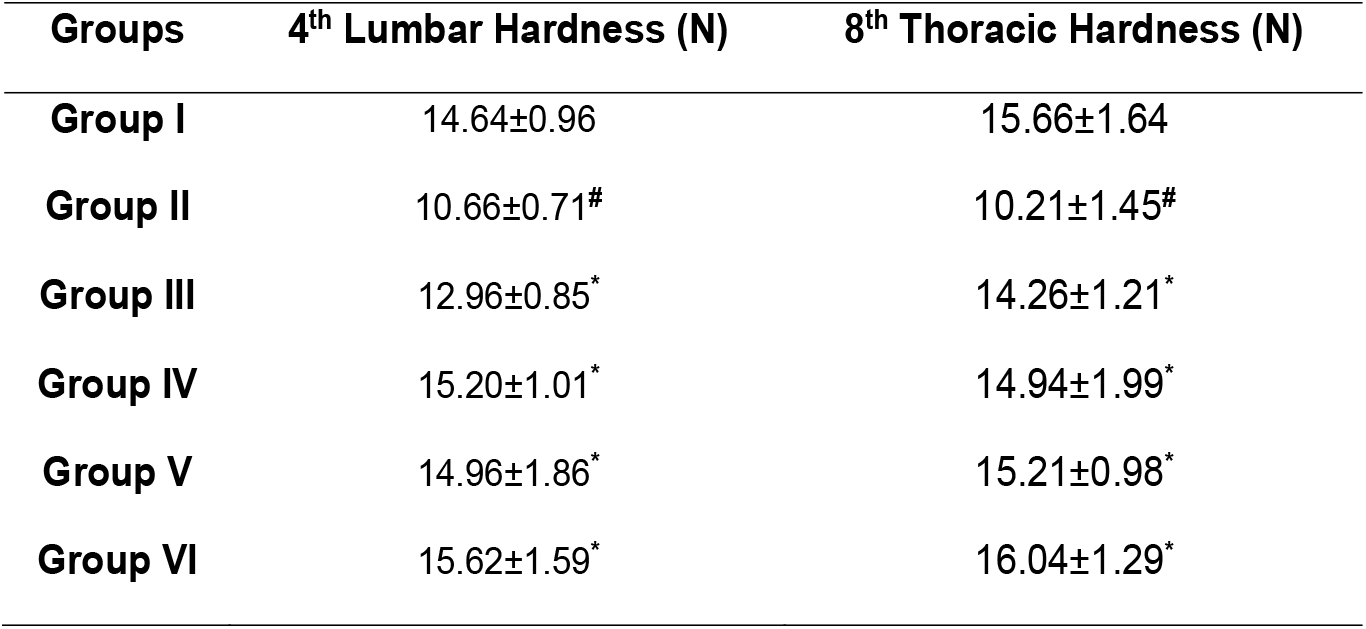
Effect of therapeutic interventions on 4^th^ lumbar vertebrae hardness (N), 8^th^ thoracic vertebrae hardness (N) [data were expressed as mean ± SD (n=6); analysed by one-way ANOVA followed by Tukey’s multiple comparison test. # = p<0.05, considered statistically significant when compared group I vs group II. * = p<0.05, considered statistically significant when compared to the other treatment groups (group III, IV, V, VI) vs. diet control group]

| Groups | 4<sup>th</sup> Lumbar Hardness (N) | 8<sup>th</sup> Thoracic Hardness (N) |
| --- | --- | --- |
| Group I | 14.64±0.96 | 15.66±1.64 |
| Group II | 10.66±0.71<sup>#</sup> | 10.21±1.45<sup>#</sup> |
| Group III | 12.96±0.85<sup>*</sup> | 14.26±1.21<sup>*</sup> |
| Group IV | 15.20±1.01<sup>*</sup> | 14.94±1.99<sup>*</sup> |
| Group V | 14.96±1.86<sup>*</sup> | 15.21±0.98<sup>*</sup> |
| Group VI | 15.62±1.59<sup>*</sup> | 16.04±1.29<sup>*</sup> |

### Effect of therapeutic interventions on femur bone mineral density (g/cm2)

There was a significant (p ◻0.05) decline in the bone mineral density of animals fed with DC animals compared to normal animals. Whereas, in case of animals treated with other therapeutic interventions (groups III, IV, V, VI) showed a significant (p ◻0.05) increase in the bone mineral density compared to animals fed with vitamin D deficient diet (Table 3).

Among all the studied interventions, group VI (chrysin + CaCO_3_ + vitamin D_3_) showed better improvement in all parameters (serum, urinary, bone).

### Effect of therapeutic interventions on 25-OH-D_3_ by LC-MS/MS

A significant (p ◻0.05) decrease in serum 25-OH-D_3_ levels of animals treated with vitamin D deficient diet was noticed compared to normal animals which correlates with the altered biochemical and bone parameters. But, comparatively there was a significant (p ◻0.05) increase in 25-OH-D_3_ levels in serum samples of animals treated with other therapeutic interventions i.e., III, IV, V, VI were observed in contrast with group II animals (Figure 1).

**Figure 1:**
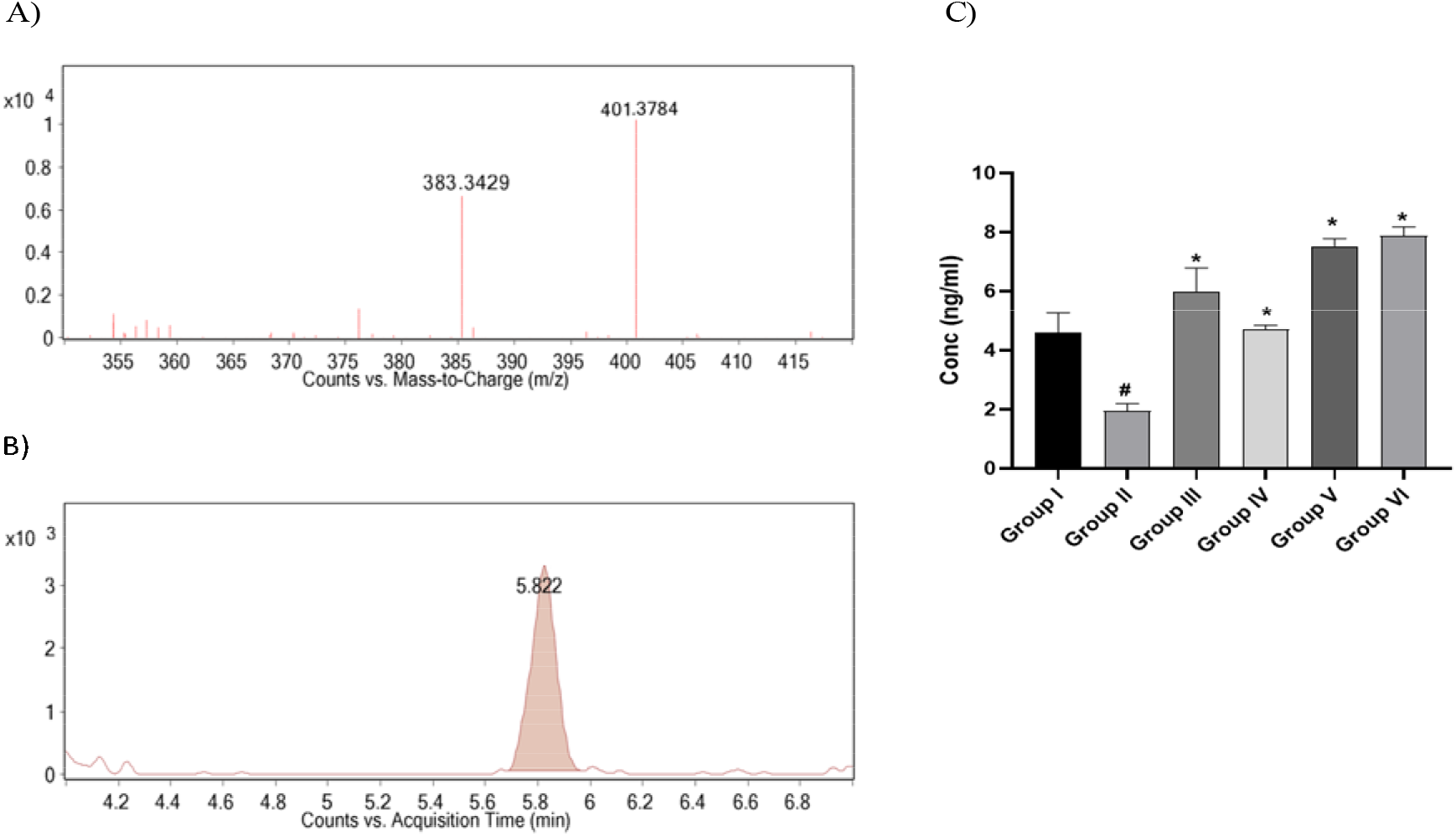
Quantification of 25-OH-D_3_ levels by LC-MS/MS. **(A)** Mass fragmentation of 25-OH-D_3_ (B) Ion chromatogram of the selected fragment ion (m/z 383.5→401.3). (C) Quantification of serum 25-OH-D_3_ from area under the curve [data were expressed as mean ± SD (n=4); analysed by one-way ANOVA followed by Tukey’s multiple comparison test. # = p<0.05, considered statistically significant when compared group I vs group II. * = p<0.05, considered statistically significant when compared to the other treatment groups (group III, IV, V, VI) vs. diet control group]

### Effect of therapeutic interventions on histopathology of femur bone

The effect of therapeutic interventions on histopathology of femur bone was studied at 10X magnification and the detailed description was mentioned, and photomicrographs were given in the figure 2.

**Figure 2:**
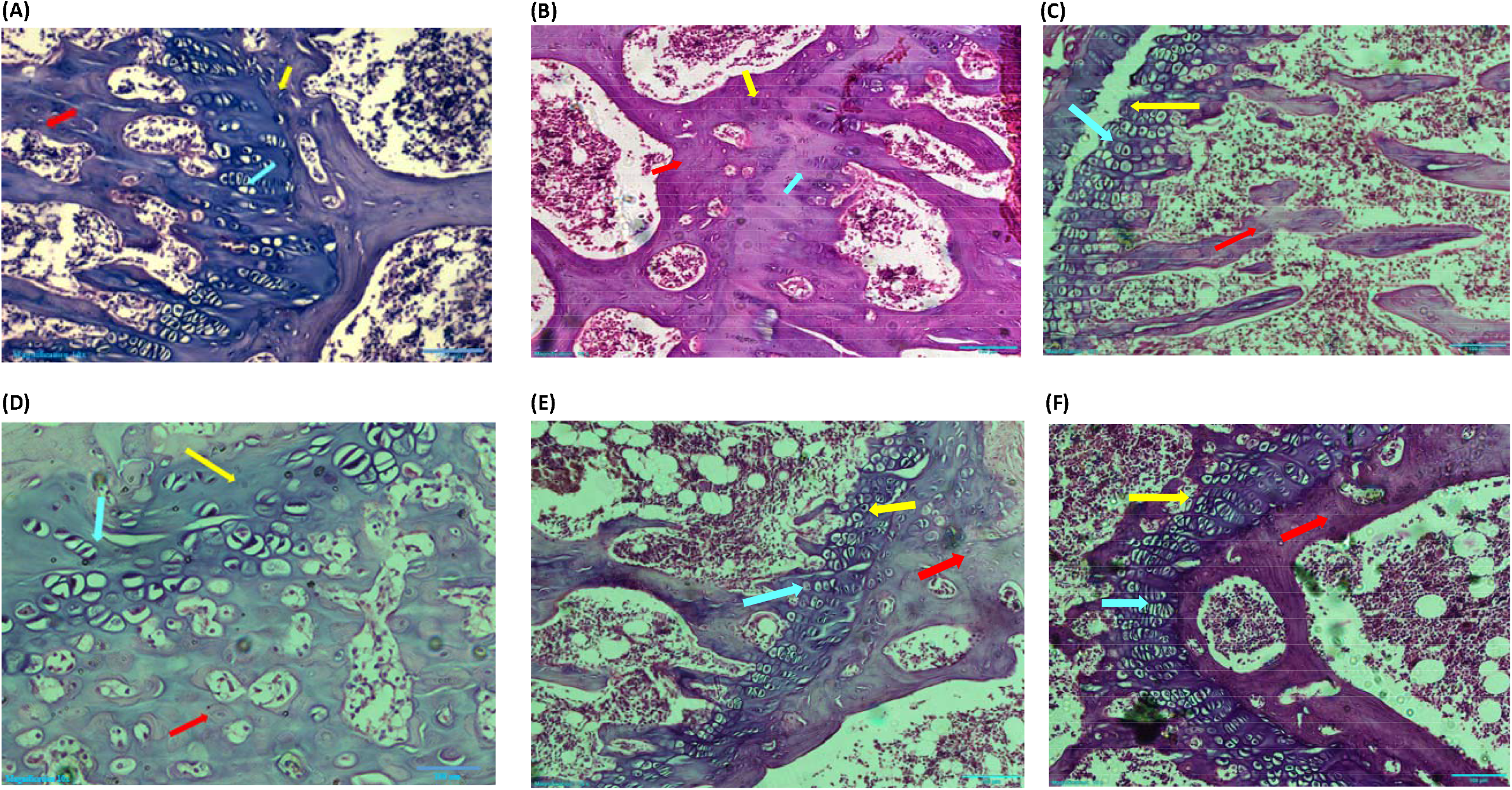
Impact of chrysin on histopathology of bone. **A)** Group I (normal group); **B)** Group II (disease control); **C)** Group III (chrysin); **D)** Group IV (chrysin + CaCO_3_); **E)** Group V (chrysin + vitamin D_3_); **F)** Group VI (chrysin + vitamin D_3_ + CaCO_3_) showing alterations in normally growing cells (blue arrow) at epiphyseal plate (yellow arrow) along with osteocytes (red arrow) in bone matrix.

The photomicrographs of the group I (normal) have shown the normal distribution of cells in the epiphyseal plate and also the even distribution of osteocytes in the bone matrix. Whereas there was a decreased incidence of differentiating osteocytes and bone formation cells in the epiphyseal region was spotted among the animals fed with group II (vitamin D deficient diet). Osteoclasts were also prominently noticed in the DC animals which supports the process of bone resorption. However, in the case of animals treated with group III (chrysin) showed number of osteocytes at the epiphyseal plate and bone formation cells such as osteoblasts were observed. In group III few osteoclasts were also spotted but not as much as compared to vitamin D deficient control group.

When it comes to group IV (chrysin + CaCO_3_) increased proliferation of cells in the epiphyseal plate was observed compared to the diet control group, followed by subsequent calcification. In group V (chrysin + vitamin D_3_) considerable even distribution of osteocytes and matrix formation was observed which indicated active bone growth. Moreover, epiphyseal plate architecture was observed to be better than the diet control group. In group VI (chrysin + CaCO_3_+ vitamin D_3_) increased bone formation due to increased osteoblast cells in the epiphyseal plate of bone and comparatively less resorptive cells in contrast with diet control group were noticed.

## Discussion

Vitamin D has gained prominence value as nutrient supplement from diverse sources and as an endogenous hormone biosynthesized from cholesterol by UV irradiation. Since from the few decades sunshine vitamin has gained an optimistic importance in various pathological conditions in addition with former osseous effects [33]. Evidence stated that appropriate levels of serum 25-OH-D_3_ has specified their essential role in homeostasis of essential nutrients such as calcium, magnesium, and phosphorus which accentuates bone mass and mineralization. Several studies were reported that alterations in these mineral nutrients would result in elevated bone turnover most significantly in vitamin D deficient condition [34].

As the females are more prone to osteoporotic fractures and bone loss female rats were preferred for the current study [35]. In our study, eminent rise in the serum calcium, magnesium, and phosphorus levels were obtained among animals treated with different therapeutic interventions suggests their increased availability accompanied by the decline in urinary excretion by which it was noticed that examined combinations of chrysin with CaCO_3_ and/or vitamin D_3_ were showing potential interactions which may also have a further impact on bone mineralization. Similar studies are available which support alteration in the serum and urinary levels of calcium with vitamin-D deficient diet and therapeutic interventions [36, 9]. But lot more studies are warranted to conclude that this progress is dependent on decreased urinary excretion or may be due to increased absorption or decreased metabolism.

In contrast, the animals fed with vitamin D deficient diet (group II) doesn’t showed any variance in calcium levels compared with normal animals which convey the diet provided doesn’t have any modulatory effect on calcium levels among DC animals. Nevertheless, diminished serum magnesium and phosphorus levels among vitamin D deficient animals assures the importance of vitamin D in maintaining appropriate mineral homeostasis. On the same vein, the DC animals showed enhanced excretion urinary calcium, phosphorus and magnesium which furthers supports the importance of vitamin D in renal excretion of essential minerals respectively [37, 38, 9].

In this context, intensified serum ALP level was noticed among group II animals suggests the induction of bone ill-health which also assert the dynamic imbalance of bone tissue promoting bone resorption which consistently supports the previous reports obtained [39]. Furthermore, a consistent decline in the serum ALP levels was obtained in animals treated with other therapeutic groups (groups III, IV, V, VI) indicating the considerable recovery of disease condition.

Another supportive data obtained from the study presented a positive correlation between body weights and bone parameters such as weight, hardness of tibia-fibula, femur of the animals with course of chrysin treatment retrieve bone mass and importance of vitamin D in bone mineral absorption. In contrary, group II animals mitigated this effect which resembles the results obtained from earlier studies indicating the increased body weight is proportional to the bone weight [40,41]. No changes in the lengths of the femur bone and tibia-fibula bone were observed.

In addition, the diminished availability of serum biochemical estimations of essential minerals such as calcium, magnesium and phosphorus were witnessed in animals with insufficient vitamin D (group II) can also be analogous with the decreased hardness in 4^th^ lumbar and 8^th^ thoracic vertebrae suggests the deterioration in bone mass that corroborates previous studies. Comparatively, other therapeutic groups showed amplified bone strength with respect to increased 4^th^ lumbar and 8^th^ thoracic vertebral hardness. A comparable effect in bone strength was observed with increased serum calcium, magnesium and phosphorus in earlier reports [28]

Previous research revealed the imperative link between the bone mineralisation and bone ash content in bone related diseases. In this regard, vitamin D deficient diet control group had shown decreased bone ash content i.e., calcium, phosphorus, magnesium which confirms decreased bone mineral deposit [42]. Nevertheless, all the treatment groups have shown enhanced calcium content in bone ash, which can be attributed to the increased serum calcium levels. None of the therapeutic interventions caused any increase in magnesium deposits in the bones. But chrysin, when given in combination with vitamin D_3_ and CaCO_3_ has caused a prominent increase in the phosphorus content in the bone ash. The role of CaCO_3_ and vitamin D in increasing bone mineralization has been established in earlier studies [43].

Femur bone mineral density (BMD) is one of the gold standard tests for the determination of mineral strength of bone was assessed by performing DEXA (30). The animals treated with vitamin D deficient diet showed decreased bone mineral density compared to the normal group which indicates decreased bone calcification, mineralisation, and bone mass. Significantly, animals in different treatment groups showed intense bone mineral density which supports the results obtained from biochemical and bone ash estimations [31].

In order to estimate the inhibitory effect of chrysin on vitamin D metabolism and also its effect on low dose vitamin D_3_ (40IU) supplementation, estimation of serum 25-OH-D_3_ levels were considered as a reliable marker for vitamin D status [44]. From our experimental analysis, the diet control group showed decreased serum 25-OH-D_3_. In contrary, increased serum 25-OH-D_3_ levels were obtained in the animals treated with different therapeutic interventions reinforces its physiological effect on bone associated with reports of serum biochemical parameters discussed above. Chrysin when given alone effectively elevated 25-OH-D_3_ levels, but the combination of chrysin with CaCO_3_ and vitamin D_3_ has shown a major impact in increasing the serum 25-OH-D_3_ levels which partly confirms the metabolic interactions of chrysin on vitamin D catabolism even in low dose of vitamin D_3_ (40IU) as well.

The histopathological data also revealed the greater number of bone formation cells in the epiphyseal plate of the animals treated with different therapeutic interventions which also favours the data obtained from other biochemical parameters. But, uneven distribution of osteocytes in the bone matrix were spotted in animals along with a smaller number of bone formation cells in the epiphyseal plate among diet control group (group II) bone specimens resembles decreased bone formation and can be attributed to hypovitaminosis D as mentioned earlier [45]. Hence, this study showed the administration of chrysin along with vitamin D_3_ and CaCO_3_ can be an effective strategy for treating bone disorders in vitamin D insufficient condition.

## Conclusion

Our preclinical research results revealed that the combination of chrysin + vitamin D_3_ + CaCO_3_ showed a greater implication on bone health than all other combinations studied. The result also clearly indicates that chrysin was able to increase availability of active serum vitamin D even with minimal vitamin D_3_ supplementation (40IU), but to confirm interactions at the site of metabolism, further studies are warranted. The study has opened a new option to treat vitamin D deficiency which can be used as a cost-effective option to improve bone health.

## Supporting information

The composition of the vitamin D deficient diet as per the reference [22] was given in table 1

## Acknowledgements

Authors acknowledge UGC-SAP, DST-FIST of Institute of Pharmaceutical Technology and DST-CURIE of Sri Padmavati Mahila Visvavidyalayam for providing the infrastructural facilities to carry out the study. The support of Dr. G. Sireesha, Department of Home Science, Sri Padmavati Mahila Visvavidyalayam is acknowledged for preparation of vitamin D deficient diet. Dr. Vasudharani Devanathan, Indian Institute of Science Education and Research, Tirupati has provided support for microscopical studies.

## Author contribution

Siva Swapna Kasarla has carried out the experimentation work. The design and supervision of the study was done by Sujatha Dodoala. The analytical work and bone mineral density estimations were carried out under the supervision of Sunitha Sampathi and Narendra Kumar Talluri.

## References

1. Briggs AM, Woolf AD, Dreinhofer K, Homb N, Hoy DG, Kopansky-Giles et al. Reducing the global burden of musculoskeletal conditions. Bulletin of the World Health Organization. 2018;96(5):366–368.

2. Hyder AA, Wosu AC, Gibson DG, Labrique AB, Ali J, Pariyo GW. Non communicable disease risk factors and mobile phones: A proposed research agenda. J Med Internet Res. 2017;19(5):1–10.

3. Pouresmaeili F, Kamalidehghan B, Kamarehei M, Goh YM. A comprehensive overview on osteoporosis and its risk factors. Ther Clinical Risk Manag. 2018;14:2029–2049.

4. Kelsey JL. Risk factors for osteoporosis and associated fractures. Public Health Rep. 1989;104:14–20.

5. Christodoulou S, Goula T, Ververidis A, Drosos G. Vitamin D and Bone Disease. Biomed Res Int. 2013;1–6.

6. Laird E, Ward M, McSorley E, Strain JJ, Wallace J. Vitamin D and Bone Health; Potential Mechanisms. Nutrients. 2010;2:693–724.

7. Dusso AS, Brown AJ, Slatopolsky E. Vitamin D. Am J Physiol Renal Physiol. 2005;289: F8–F28.

8. Khazai N, Judd SE, Tangpricha V. Calcium and vitamin D: Skeletal and extraskeletal health. Curr Rheumatol Rep, 2008;10:110–117.

9. Nair P, Venkatesh B, Center JR. Vitamin D deficiency and supplementation in critical illness-the known knowns and known unknowns. Crit Care. 2018;22:3–9.

10. Abulmeaty MMA. Sunlight exposure vs. vitamin D supplementation on bone homeostasis of vitamin D deficient rats. Clin Nutr Exp. 2017;11:1–9.

11. Silva MC, Furlanetto TW. Intestinal absorption of vitamin D: a systematic review. Nutr Rev. 2017;76(1):60–76.

12. Mohseni H, Hosseini SA, Amani R, Ekrami A, Ahmadzadeh A, Latifi SM. Circulating 25-Hydroxy Vitamin D Relative to Vitamin D Receptor Polymorphism after Vitamin D3 Supplementation in Breast Cancer Women: A Randomized, Double-Blind Controlled Clinical Trial. Asian Pac J Cancer Prev. 2017;18(7):1953–1959.

13. Wikvall K. Cytochrome P450 enzymes in the bioactivation of vitamin D to its hormonal form (Review). Int J Mol Med. 2001;7:201–209.

14. Wang Z, Lin YS, Zheng XE, Senn T, Hashizume T, Scian M, et al. An Inducible Cytochrome P450 3A4-Dependent Vitamin D Catabolic Pathway. Mol Pharmacol, 2011;81(4):498–509.

15. Wang Z, Schuetz EG, Xu Y, Thummel KE. Interplay between vitamin D and the drug metabolizing enzyme CYP3A4. J Steroid Biochem Mol Biol. 2013;136:54–58.

16. Bibi Z. Role of cytochrome P450 in drug interactions. Nutr Metab (Lond). 2014;11(1):11.

17. Kearns MD, Alvarez JA, Tangpricha V. Large, Single-Dose, Oral Vitamin D Supplementation in Adult Populations: A Systematic Review. Endocr Pract. 2014;20(4):341–351.

18. Sergent T, Dupont I, Vander Heiden E, Seippo ML, Pussemier L, Larondelle Y et al. CYP1A1 and CYP3A4 modulation by dietary flavonoids in human intestinal caco – 2 cells. Toxicol Lett. 2009;191;216–222.

19. Mani R, Natesan V. Chrysin: Sources, beneficial pharmacological activities, and molecular mechanism of action. Phytochemistry. 2018;145:187–196.

20. Kaczmarczyk-Sedlak I, Wojnar W, Zych Maria, Ozimina-Kamińska, Sedlak L. Effect of chrysin on the mechanical properties of bones in ovariectomized rats. In: Bogucka-Kocka A, Kocki J, Sowa I. Proceedings of the 4^th^ International Conference on Plant-The Source of Research Material; 2015 Sep; and Workshop. Lublin, Poland. 129.

21. Coman C, Vlase E. Formulation, preparation and chemical analysis of purified diets for laboratory mice and rats. Scientific Works. Series C. Vet Med. 2017;13(1):149–154.

22. Mallya SM, Corrado KR, Saria EA, Yuan FF, Tran HQ, Saucier K et al. Modeling vitamin D insufficiency and moderate deficiency in adult mice via dietary cholecalciferol restriction. Endocr Res. 2016;41(4):290–299.

23. Kouhnavard M, Esfahani NE, Montazeri M, Hashemian SJ, Mehrazma M, Larijani B, et al. Effects of Vitamin D and Calcium Supplementation on Micro-architectural and Densitometric Changes of Rat Femur in a Microgravity Simulator Model. Iran Red Crescent Med J. 2014;16(6):1–6.

24. Orsolic N, Goluza E, Dikic D, Lisicic D, Sasilo K, Rodak E, et al. Role of flavonoids on oxidative stress and mineral contents in the retinoic acid-induced bone loss model of rat. Eur J Nutr. 2014;53(5):1217–27.

25. Hardik KS, Jalak MK, Deepti KJ, Ghandshyam RP. Pharmacological investigation of Bonton capsule for Anti-osteoporotic activity in ovariectomized rat. International Journal of Pharmaceutical and Phyto-pharmaceutical research. 2013;3:52–56.

26. Adamec J, Jannasch A, Huang J, Hohman E, Fleet JC, Peacock M, et al. Development and optimization of an LC-MS/MS-based method for simultaneous quantification of vitamin D2, vitamin D3, 25-hydroxyvitamin D2 and 25-hydroxyvitamin D3. J Sep Sci. 2010;34(1):11–20.

27. Zhang SW, Jian W, Sullivan S, Sankaran B, Edom RW, Weng N, et al. Development and validation of an LC–MS/MS based method for quantification of 25 hydroxyvitamin D2 and 25 hydroxyvitamin D3 in human serum and plasma. J Chromatogr B Analyt Technol Biomed Life Sci. 2014;961:62–70.

28. Halekunche Y, Burdipad G, Kuppusamy S, et al. Anti–osteoporotic activity of ethanol extract of leave on Punica granatum leaves on ovariectomized rats. Asian J Pharm Pharmacol. 2016;2(4):85–92.

29. Chitme HR, Muchandi IS, Burli SC. Effect of Asparagus Racemosus Wild Root Extract on Ovariectomized Rats. The Open Natural Products Journal. 2009;2:16–23.

30. Lu PW, Briody JN, Howman-Giles R, Trube A, Cowell CT. DXA for bone density measurement in small rats weighing 150–250 grams. Bone. 1994;15(2):199–202.

31. Lucinda LMF, Aarestrup BJV, Reboredo MM, Pains TDA, Chaves ZR, Chaves RZ, et al. Evaluation of the anti-osteoporotic effect of Ginkgo biloba L. in Wistar rats with glucocorticoid-induced-osteoporosis by bone densitometry using dual-energy x-ray absorptiometry (DEXA) and mechanical testing. An Acad Bras Cienc. 2017;89(4): 2833–2841.

32. Shirwaikar A, Khan S, Malini S. Antiosteoporotic effect of ethanol extract of Cissus quadrangularis Linn. on ovariectomized rat. J Ethnopharmacol. 2003;89(2-3): 245–250.

33. Zhang R, Naughton DP. Vitamin D in health and disease: Current perspectives. Nutr J. 2010;9(1):1–13.

34. Schwalfenberg GK, Genuis SJ. Vitamin D, Essential Minerals, and Toxic Elements: Exploring Interactions between Nutrients and Toxicants in Clinical Medicine. ScientificWorldJournal. 2015;1–8.

35. Alswat KA. Gender Disparities in Osteoporosis. J Clin Med Res. 2017;9(5):382–387.

36. Toromanoff A, Ammann P, Mosekilde L, Thomsen JS, Riond JL. Parathyroid hormone increases bone formation and improves mineral balance in vitamin D-deficient female rats. Endocrinol. 1997;138(6):2449–57.

37. Yamamoto M, Kawanobe Y, Takahashi H, Shimazawa E, Kimura S, Ogata E. Vitamin D deficiency and renal calcium transport in the rat. J Clin Invest. 1984;74(2):507–13.

38. Keenan MJ, Hegsted M, Siver F, Mohan R, Wozniak P. Recovery of Rats from Vitamin D-Deficient Mothers. Ann Nutr Metab. 1991;35(6):315–327.

39. Grigoryan AV, Dimitrova AA, Kostov KG, Russeva AL. Atanasova MA, Blagev AB, et al. Changes of Serum Concentrations of Alkaline Phosphatase and Metalloproteinase-9 in an Ovariectomized Wistar Rat Model of Osteoporosis. J Biomed Clin Res. 2017;10(1):32–36.

40. Yang X, Li F, Yang Y, Shen J, Zou R, Zhu P, Zhang C, et al. Efficacy and Safety of Echinacoside in a Rat Osteopenia Model. Evid Based Complement Alternat Med. 2013:1–10.

41. Wang C, Zhang Y, Xiong, Y, Lee CJ. Bone composition and strength of female rats subjected to different rates of weight reduction. Nutr Res. 2000;20:1613–1622.

42. Mustafa RA, Alfky NAA, Hijazi HH, Header EA, Azzeh FS. Biological effect of calcium and vitamin D dietary supplements against osteoporosis in ovariectomized rats. Prog Nutr. 2018;20(1):86–93.

43. Khazai N, Judd SE, Tangpricha V. Calcium and vitamin D: skeletal and extraskeletal health. Curr Rheumatol Rep. 2008;10(2):110–117.

44. Holick MF. Vitamin D Status: Measurement, Interpretation, and Clinical Application. Ann Epidemiol. 2009:19(2);73–78.

45. Fossey S, Vahle J, Long P, et al. Nonproliferative and Proliferative Lesions of the Rat and Mouse Skeletal Tissues (Bones, Joints, and Teeth). J Toxicol Pathol. 2016;29(3 Suppl):49S–103S.

