## Supplementary figures and images for "Impact of Chrysin on Vitamin D and Bone Health - Preclinical Studies"

**Table 1: The composition of the vitamin D deficient diet as per the reference [22].**

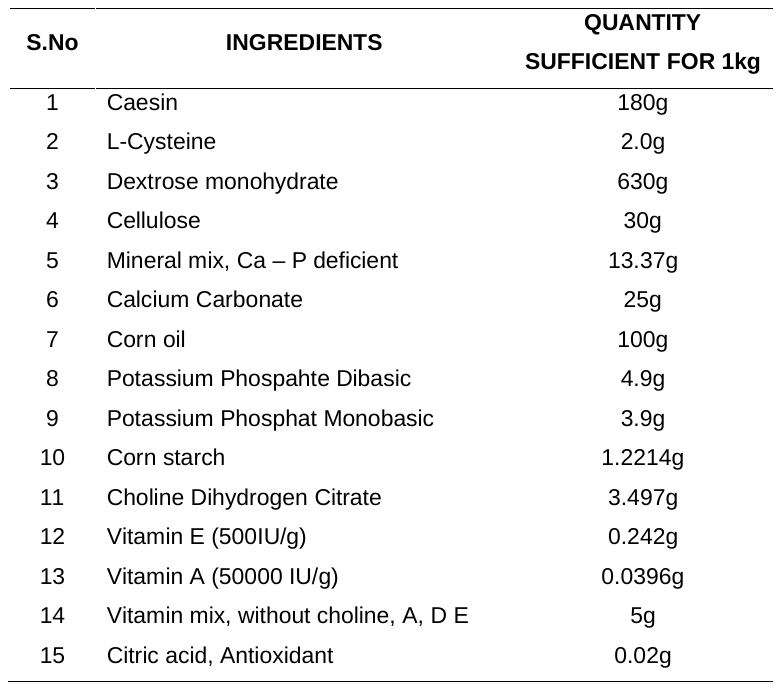

Supplement: The composition of the vitamin D deficient diet as per the reference [22] was given in table 1 [file 390757_file04.docx]
